# Frequency tagging evidence supports perceptual separation of rapid stimuli in human fetuses

**DOI:** 10.1101/2025.06.06.658307

**Authors:** Joel Frohlich, Julia Moser, Dimitrios Metaxas, Katrin Sippel, Laura Batterink, Hubert Preissl

## Abstract

In early human development, perceptual processes grow faster with maturation, as inferred using the duration of the attentional blink and multisensory integration window. The consequences of this developmental trend for sensory-cognition in fetuses are unclear: does the fetus perceive rapid stimuli as discrete events or, rather, one fused stimulus? We addressed this question using frequency tagging in two experiments with rapid auditory stimuli while neural responses were recorded in the third trimester with fetal magnetoencephalography (MEG). Our results are the first successful demonstration of frequency tagging in the fetal MEG amplitude spectrum and show that the fetal cortex generates separate neural responses to discrete auditory stimuli in both experiments, and similar results were also obtained when one experiment was repeated in newborns. While we cannot rule out perceptual fusion of rapid stimuli in higher-order association cortices, our results weaken the hypothesis that fetuses fuse rapid auditory stimuli into a single prolonged percept. Finally, our work points to frequency tagging analysis as a solution which avoids the uncertainties surrounding immature neural response latencies in time-domain analysis of fetal MEG.

## Introduction

As the brain develops, so too does the pace of neural information processing and the temporal grain of perception, likely as a result of increasing myelination (1). For instance, evidence from developmental studies, including work in human infants and children, indicates that both the attentional blink (2) and the multisensory integration (3) window grow shorter with age, corresponding to faster cognitive-perceptual processing. Because most myelination occurs after birth (4), one might expect that the speed of perceptual processing is severely limited before birth. Indeed, extrapolating from the developmental trend above, the temporal grain of fetal awareness might be too coarse for rapid stimuli (i.e., repetition frequency *≥* 1 Hz) to be perceived by the fetus as discrete events. Addressing this possibility carries implications for the design of experiments that probe prenatal perception, attention, awareness, and working memory (5, 6). Recent years have seen increasing interest in the womb as a last frontier of cognitive and consciousness sciences (7–15), highlighting the timeliness of this problem.

Herein, we tested the hypothesis that rapid stimuli are perceptually smeared or fused by the fetus into a prolonged whole (henceforth: “fused stimulus hypothesis”) using frequency tagging with fetal magnetoencephalography (MEG) recorded during rapid auditory stimulation in the third trimester of pregnancy. Fetal MEG is a non-invasive tool for recording fetal cortical signals after 25 weeks gestational age using superconducting magnetometers (10). While a previous fetal MEG study (16) examined evoked field responses to stimuli with a 4.5 Hz repetition frequency, frequency tagging was not attempted. A later fetal MEG pilot study (17) demonstrated neural steady state responses to amplitude modulated auditory stimuli, yet these stimuli were continuous carrier waves rather than trains of discrete stimuli. More recently, an electroencephalography (EEG) study of preterm infants found frequency-tagging in response to discrete, rapid stimuli (18), yet preterm infants are questionable models of age-equivalent fetuses due to differences in health and, moreover, environment (9); the process of birth itself likely triggers many developmental processes relevant for sensory-driven cognition (9, 19). Thus, no study to date has adequately addressed whether the fetal brain can accurately resolve rapid auditory stimuli.

We asked whether the fetal brain resolves subsecond inter-stimulus intervals. Specifically, we investigated this question with two fetal MEG experiments. In Experiment 1, structured and pseudo-randomly ordered auditory tones were played continuously at a rate of 3 tones per second (3 Hz). In Experiment 2, fetuses were exposed to four-tone auditory sequences at a repetition frequency of 1.67 Hz in the context of a local-global oddball paradigm (5, 6). A subset of subjects in Experiment 2 then returned to the laboratory after birth to participate in the same passive auditory paradigm again as newborn infants. Both experiments yielded positive results at the group level, thus showing, for the first time, that the signal-to-noise ratio (SNR) of the group-averaged fetal MEG amplitude spectrum is sufficient to detect frequency tagging in the developing cortex.

## Methods

Both experiments were approved by the local ethics committee of the Medical Faculty of the University of Tuebingen and were conducted in accordance with the Declaration of Helsinki. Written informed consent was obtained from the mother-to-be (fetuses) or both parents (newborns). MEG data were recorded in a magnetically shielded room at the University of Tuebingen fMEG Center. After strict quality control, Experiment 1 yielded 31 recordings from 31 fetuses aged 31 - 38 gestational weeks. Experiment 2 yielded 81 recordings, including longitudinal data in which fetuses were recorded from at multiple gestational ages, from 43 fetuses aged 25 - 40 gestational weeks, as well as 20 recordings from 20 newborns with full-term births aged 40 - 49 gestational weeks at the time of data collection. The same MEG device was used to record from both fetuses and newborns. Stimuli are described in previous publications by Moser et al. [Experiment 1: (20); Experiment 2: (5, 21)].

MEG signals were bandpass filtered in different frequency ranges for fetuses and newborns, accounting for the greater noise inherent in the former (fetuses: 1 - 10 Hz; newborns: 1 - 15 Hz), and averaged across trials and channels following in-house preprocessing pipelines (5, 21–23); see Supplemental Methods for details. Data analyses were carried out in MATLAB R2022a and R2025a using all usable data following channel and trial-averaging. Because each MEG recording encompassed multiple stimulus conditions which all shared a common stimulation frequency, conditions were concatenated across the dimension of time; see Supplemental Methods for details. For Experiment 1, we limited our analysis to the first 6 minutes of data in each of two conditions (cf. *≤* 5 minutes of data in prior infant EEG frequency tagging work by de Heering and Rossion (24)), as we reasoned that data in the second half of the experiment were more likely to induce habituation (25), because the stimuli were a constant stream of regularly spaced auditory tones. For Experiment 2, all usable trials from the test phase were averaged for each condition, as stimuli were grouped into sequences punctuated by silent pauses rather than a constant stream, making habituation less likely.

Next, to enhance SNR, we computed the group-averaged signal across all fetal and neonatal recordings, separately, in each experiment. To test for a frequency tagging effect at the repetition frequency of the auditory stimuli (Experiment 1: 3.0 Hz; Experiment 2: 1.67 Hz), we computed the fast Fourier transform (FFT) of each group-averaged signal and then computed the SNR at the frequency of interest from the FFT amplitude spectrum referenced to adjacent frequency bins as a noise estimate, as per prior studies of frequency tagging in infant EEG (24, 26, 27). The SNR was converted to a z-score using the mean and standard deviation across adjacent frequency bins. For parametric hypothesis testing, this z-score was converted to a one-tailed P-value using a Z-test.

In addition to parametric testing, we introduced a nonparametric testing method for evaluating frequency tagging which, unlike a Z-test, makes no assumptions about the data distribution. To confirm its reliability, the nonparametric approach was validated on a publicly available dataset of infant EEG frequency tagging data (24). For nonparametric hypothesis testing, we generated a null distribution of SNR by creating surrogate data from each recording prior to group-averaging, thus destroying temporal alignment of neural events between recordings. Surrogate data were generated using the IAFFT1 algorithm by Shreiber et al. (28), which, unlike common FFT phase randomization procedures (29), preserves both the signal’s power spectrum and its exact data distribution. This procedure was repeated 1000 times, and an empirical P-value was computed by comparing the actual SNR with the dull distribution.

Finally, to ensure that our results were specific to the hypothesized frequency, we repeated each analysis with the frequency of interest swapped between experiments as a control (i.e., control frequency in Experiment 1: 1.67 Hz; Experiment 2: 3.0 Hz). To correct for multiple testing, we applied the false discovery rate (FDR) procedure by Benjamini and Hochburg (30) to all P-values; this was done separately for experimental and control results, as well as for the main results (Table 1) and supplemental results (Table S1).

**Table 1.**
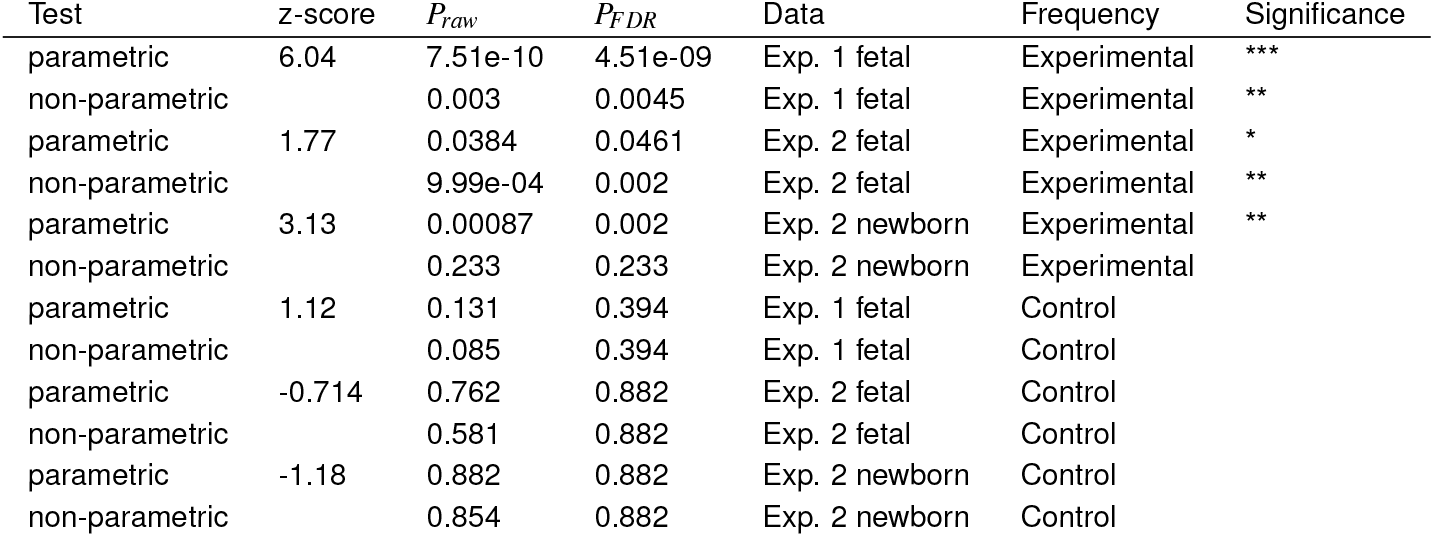
Statistical analysis of frequency tagging strength in each experiment. (Exp. = experiment) *P*_*FDR*_ < **0.05** * *P*_*FDR*_ < **0.005** ** *P*_*FDR*_ < **0.0005** ***

For detailed methods, including supplemental analyses, please see the Supplemental Material.

## Results and Discussion

### Evidence of MEG frequency tagging in fetuses and newborns

We applied both parametric and nonparametric approaches to MEG data from fetuses and newborns to investigate the timescale of fetal perceptual processing. At the group level, each fetal or newborn dataset in each experiment showed strongly significant frequency tagging (*P*_*FDR*_ < 0.005) according to at least one of the two approaches (Table 1; Fig. 1). Conversely, a significant frequency tagging effect was never observed when a control frequency was tested (Table S1, all adjusted P-values converged to *P*_*FDR*_ = 0.88). The positive results obtained at the experimental frequency are unlikely to be the result of cardiac artifacts, as the peaks in the amplitude spectra we obtained from group-averaged MEG signals (Fig. 1A,C,D) do not demonstrate the harmonics characteristic of the fetal or maternal MCG signal amplitude spectra (Fig. S1). To confirm that our newly introduced nonparametric testing approach is sensitive to frequency tagging effects, we showed similar results using both parametric and nonparametric approaches in publicly available infant EEG frequency tagging data (24) (Fig. S2).

**Figure 1.**
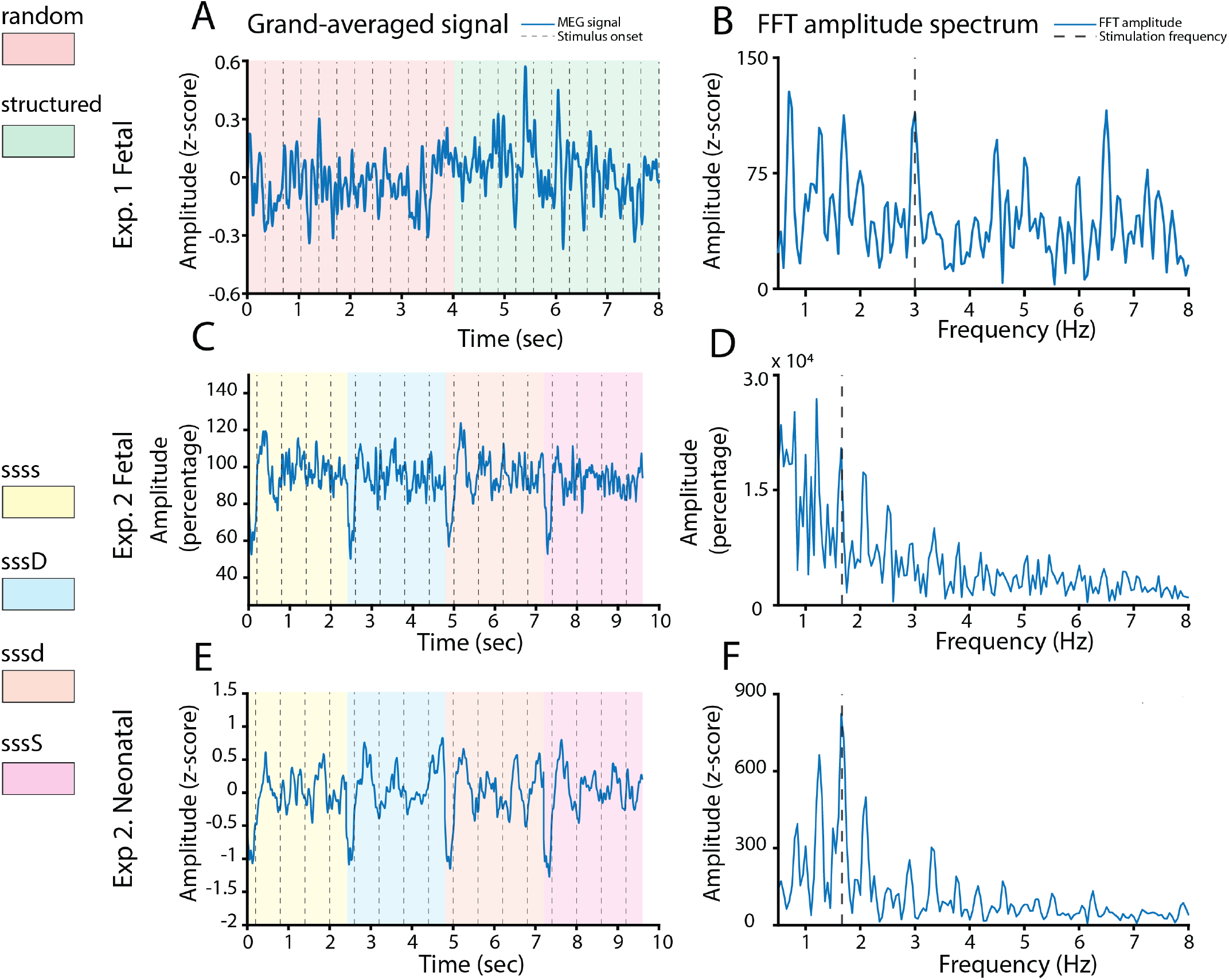
Convergent evidence against the fused stimulus hypothesis was obtained from two experiments. Experiment 1 (A,B) examined auditory stimuli with a 3.0 Hz repetition frequency in fetuses, where the MEG signal was normalized using z-scoring. Experiment 2 examined auditory stimuli with a 1.67 Hz repetition frequency in fetuses (C,D) — where the MEG signal was normalized with respect to the fetal neural response 50 - 350 ms after the first tone of the stimulus sequence — and newborns (E,F) — where the MEG signal was normalized using z-scoring. Group averaged signals are shown in the first column of panels (A, C, E), and the onset of each stimulus is marked with a vertical dashed line. FFT amplitude spectra are shown in the second column on panels (B, D, F), and stimulation frequency is marked with a vertical dashed line. The legend in the left margin shows the stimulus conditions which were concatenated to create the group-averaged signals; see Supplemental Methods for details.

Our findings suggest that the fetal brain resolves subsecond interstimulus intervals. Furthermore, the results are a technical accomplishment insofar as they show the SNR of fetal MEG is sufficient to demonstrate this effect using frequency tagging. To facilitate future independent replication, we performed a subsampling-based power analysis of Experiment 1, given that this experiment had no longitudinal fetal data and thus allowed us to treat each fetus as an independent sample. Using this approach, we determined that n = 23 fetuses are needed to detect a statistically significant frequency tagging effect with 80% statistical power using a Z-test (Fig. S3). While the amplitude spectrum was always computed from the FFT without windowing or smoothing for analysis purposes, readers interested in seeing the smoothed power spectrum should refer to Fig. S4. As a further supplemental analysis (Table S1), we used bootstrapped resampling (Fig. S5) to estimate correlations between frequency tagging strength and several perinatal variables: gestational age, birth weight, maternal body mass index or BMI, maternal age, fetal/neonatal sex, and fetal/neonatal heart rate variability. We found that perinatal variables did not consistently correlate with frequency tagging strength (see Supplemental Results).

Our study benefited from a combination of both parametric statistical testing, which is consistent with prior infant EEG frequency tagging work (24, 27), and nonparametric statistical testing, which addresses the fact that the Z-test employed by the parametric approach assumes a normal distribution. The nonparametric approach tests against a very large possibility space formed by surrogate data and may therefore be less sensitive to peaks that are small on a global scale. Experiment 1 yielded a locally prominent 3 Hz amplitude peak (Fig. 1) that resulted in a strongly significant (*P*_*FDR*_ = 0.00032) parametric result, whereas the same peak was less strongly significant (*P*_*FDR*_ = 0.022) when measured nonparametrically. Conversely, fetal and neonatal results from Experiment 2 showed the opposite pattern with respect to parametric and nonparametric testing (Table 1), possibly owing to amplitude peaks that were less locally prominent than that in Experiment 1 (parametric test), yet stronger on a global scale spanned by a large sampling of surrogate data (nonparametric test); see Fig. 1. In this sense, it appears that different stimulation paradigms yield different frequency tagging signatures, even after differences in the stimulation frequency are acknowledged (Table 1). Besides differences in stimulation frequency between experiments, differences in the two stimulation paradigms may have also resulted in distinct frequency tagging signatures. For instance, fetal auditory habituation — previously demonstrated in other work (25) —-may have been rapidly induced by the continuous stream of 3 Hz stimulation used in Experiment 1, resulting in a locally prominent yet overall smaller amplitude peak (Fig. 1B). By contrast, Experiment 2 delivered sequences of auditory tones as trials separated by silent intervals longer than 1 second; thus, this experimental design may have better resisted habituation, resulting in a different frequency profile (Fig. 1D,F). While we were careful to use data from only the first six minutes of each condition in Experiment 1 for this reason, some degree of auditory habituation may have been unavoidable.

### Frequency tagging results inform estimates of temporal processing speed in fetuses

This is the first ever demonstration of fetal MEG amplitude spectrum frequency tagging. Given the evidently coarse temporal grain of human perception in infancy (2, 3, 9), the fact that frequency tagging can be demonstrated in human fetuses at frequencies > 1 Hz is somewhat surprising. Recently, for instance, 2D-ultrasound recordings failed to detect evidence of increased fetal salience, measured as looking preference, toward a 1 Hz flashing stimulus projected into the womb during the third trimester (31) — contrary to the authors’ hypothesis, the 1 Hz flashing stimulus even had significantly *diminished* salience as compared with a constant light. A provisional hypothesis that the fetal brain has difficulty perceiving rapid stimuli could perhaps make sense of the foregoing result. However, evidence for fetal MEG frequency tagging in our work challenges this hypothesis and may imply a relatively fine-grained temporal scale for prenatal perceptual processes (at least at the level of primary sensory cortices), investigations of which are highly challenging. Non-invasive electrophysiology studies in human fetuses are rare due to the high electrical resistance of the vernix caseosa, a waxy substance which protects the skin of the fetus. Until the rupture of the amniotic membranes (32), fetal neural activity can only be recorded with high temporal resolution using specialized MEG equipment (10). While fetal brain activity is more commonly recorded with functional MRI (fMRI) (33), this technique is not suited to address the fused stimulus hypothesis, as it lacks adequate subsecond temporal resolution due, in part, to the slow hemodynamic response. Fetal fMRI is also ill-suited to the question at hand because the hemodynamic deconvolution is especially challenging for fetal data, i.e., neurovascular coupling in the perinatal brain appears difficult to model (34).

Notably, a prior fetal MEG pilot study (17) has demonstrated evidence of a 27 Hz auditory steady state response to amplitude modulation of a 500 Hz carrier signal, thus laying the groundwork for our current study by suggesting that the fetal brain is sensitive to rapidly fluctuating stimulus properties. However, this work did not test whether the fetal brain could discriminate rapid trains of *discrete* stimuli, and it only provided evidence of a steady state response based on fetal MEG phase coherence. Indeed, the authors of the foregoing work (17) did not find evidence for a steady state response in the amplitude spectrum and speculated that the SNR of fetal MEG could be far too weak for amplitude-based detection. Our study builds on this earlier work using group-averaging of fetal MEG to overcome SNR limitations in data from two experiments with discrete auditory stimuli.

### Implications for fetal perception and awareness

Previously published analyses of Experiment 2 suggest that fetuses perceive deviations between — but perhaps not within — rapid auditory sequences (5). This is puzzling given that deviations between sequences are often thought *more* difficult to detect, as they require a longer memory trace in comparison to deviants within sequences. A recent review by Dehaene-Lambertz (35) introduced the view, which we refer to as the fused stimulus hypothesis, to interpret these results, speculating that stimuli with onsets separated by 600 ms might be perceived by the fetus as a single stimulus due to the ostensibly coarse temporal scale of fetal perception; see also Frohlich and Bayne (15) for a brief overview. This view would be supported if the fetal brain failed to respond to individual tones within a rapid stimulus sequence, as such evoked responses are necessary, albeit not sufficient, for the perception of individual stimuli.

By testing the fused stimulus hypothesis in the very same data that necessitated it, we showed that neural responses to individual rapid stimuli are detectable not only during the neonatal period but, indeed, even before birth. As such, our credence in the fused stimulus hypothesis is reduced, and it is plausible that fetal neural responses to auditory deviants between stimulus sequences described in a previous analysis of Experiment 2 (5) are indeed global prediction errors suggestive of prenatal consciousness (15). Nonetheless, our frequency tagging results could still be consistent with the fused stimulus hypothesis if auditory inputs are temporally fused at higher levels of cortical processing, e.g., in association cortices. Unfortunately, the low SNR and uncertainty in head position inherent to fetal MEG preclude source localization to address this possibility. Thus, at present, we cannot entirely rule out that stimuli are later fused at the level of association cortices, nor can we know with certainty how — or even if (9) — the fetus consciously perceives the stimuli in our experiments.

### Conclusions, limitations, and future directions

Our results cast doubt on the fused stimulus hypothesis while speaking to the feasibility of frequency tagging experiments with fetal MEG. Crucially, these conclusions resist several skeptical interpretations. For instance, while the 1.67 Hz amplitude spectrum peak in fetal data from Experiment 2 is close to the low frequency cutoff of the 1.0 - 10 Hz bandpass filter (Fig. 1E), this effect is unlikely to be a filtering artifact, as data from Experiment 1, which used a different stimulation frequency, did not exhibit an effect at 1.67 Hz. Furthermore, peaks observed in the amplitude spectrum are unlikely to result from cardiac artifacts, given the absence of harmonics which characterize spectra from the latter (Fig. S1). The possibility that the frequency tagging effect was generated by technical artifacts, i.e., mechanical forces that may have pushed the maternal abdomen farther from the MEG sensor array at regular intervals each time the sound balloon vibrated, is also resisted by at least two lines of evidence: 1) no such artifacts appear in the group-averaged time domain signals (Fig. 1A,D,G), despite the fact that they would be perfectly aligned with the stimulus triggers and thus easily visible, and 2) a frequency tagging effect was also present in newborns, where the auditory stimuli where presented unilaterally via an infant-friendly headphone placed contralaterally from the side of the head resting on the MEG sensor array. A shortcoming of our work, however, is that frequency tagging could not be measured at the level of individual subjects due to the low SNR of fetal MEG. As a solution, we used group-averaging, a frequency tagging approach established in previous infant (24, 27) and even adult (36, 37) EEG studies.

Our demonstration of frequency tagging in fetal MEG opens the door for frequency-domain analyses of fetal evoked responses that would be difficult to analyze in the time domain, as the evoked response latencies are unknown in fetal brain and, moreover, likely to change rapidly with each week of development in the third trimester (16). Although independent replication of our results is challenging at present due to the scarcity of bespoke hardware for fetal MEG (10), this problem will likely be solved in the near future by the proliferation of portable, non-cryogenic optically pumped magnetometers (38) which can be adopted for fetal MEG (39).

## Acknowledgments

We are grateful to all volunteers and families who participated in our research. In particular, we are especially grateful to the University of Tuebingen Women’s Clinic for their role in recruitment of pregnant participants. Furthermore, we are grateful to Pedro A. M. Mediano for guidance with the bootstrapped correlation analysis in this manuscript’s Supplement, to Tim Bayne for many enlightening discussions regarding the fused stimulus hypothesis, and to Ghislaine Dehaene-Lambertz for giving helpful comments on a presentation of this work at an infant consciousness meeting held at New York University. Finally, we thank and acknowledge funding from the FET Open Luminous project (H2020 FETOPEN-2014-2015-RIA under agreement No. 686764) as part of the European Union’s Horizon 2020 research and 2014 – 2018 training program, as well as partial funding from the Deutsche Forschungsgemeinschaft (DFG, German Research Foundation; 493345456), a grant (01GI0925) from the Federal Ministry of Education and Research (BMBF) to the German Center for Diabetes Research (DZD), and ANR-DFG French-German collaboration (PR 496/11-1).

## Author Contributions Statement

Conceptualization: J.F., J.M., H.P

Methodology: J.M., L.B., K.S.

Software/Code: J.F., J.M., K.S.

Formal Analysis: J.F., D.M.

Investigation: J.M., K.S.

Data Curation: J.M., K.S.

Original draft: J.F.

Review and editing: All authors

Visualization: J.F.

Supervision: H.P

Project Administration: H.P.

Funding Acquisition: H.P.

## License

This work is licensed under a [Creative Commons Attribution 4.0 International License](https://creativecommons.org/licenses/by/4.0/). 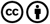

