## Supplemental Material for "Frequency tagging evidence supports perceptual separation of rapid stimuli in human fetuses"

**Joel Frohlich[1,2], Julia Moser[1,3], Dimitrios Metaxas[1,4], Katrin Sippel[1], Laura Batterink[5,6], and Hubert Preissl[1,7,8]**

[1]IDM/fMEG Center of the Helmholtz Center Munich at the University of Tübingen, Eberhard-Karls-University Tübingen, German Center for Diabetes Research (DZD), Tübingen, Germany; [2]Institute for Advanced Consciousness Studies, Santa Monica, CA, USA; [3]Masonic Institute for the Developing Brain, University of Minnesota, Minneapolis, MN, USA; [4]Graduate Training Centre of Neuroscience, University of Tübingen, Tübingen, Germany; [5]Department of Psychology, University of Western Ontario, London, Canada; [6]Centre for Brain and Mind, University of Western Ontario, London, Canada; [7]Division of Endocrinology, Diabetology and Nephrology, Department of Internal Medicine IV, University Hospital of Eberhard-Karls-University Tübingen, Tübingen, Germany; [8]German Center for Mental Health (DZPG), Partner Site Tübingen, Tübingen, Germany

This manuscript was written in LaTeX and compiled on June 6, 2025

### Supplemental Methods

#### Experiment 1

**Study population.** Experiment 1 was conducted with the original aim of investigating statistical learning in fetuses. Recruitment of pregnant women in the third trimester was undertaken by the Department of Obstetrics and Gynecology and via advertisements, e.g., announcements and emails. Inclusion criteria were as follows: singleton pregnancy without complications, appropriate fetal growth for gestational age, and full legal age (18 years or older) for the pregnant woman. Exclusion criteria were as follows: smoking, alcohol, or drug consumption during pregnancy, or other pregnancy risk factors. All prospective participants were informed, both in writing and orally, about the aims, benefit, duration and possible risks of the study. Fetal MEG signals were recorded from 37 subjects prior to excluding noisy recordings.

**Experimental details.** Following informed consent, a midwife carried out ultrasonic measurements (Ultrasound Logiq 500MD, GE, UK) to determine the fetal head position. Next, fetal magnetoencephalography (MEG) signals were recorded using the SQUID Array for Reproductive Assessment (SARA; SQUID = superconducting quantum interference device) in a magnetically shielded room (Vakuumschmelze, Germany). The fetus was exposed to two blocks of auditory tone sequences — one structured and one random — during the MEG recording. In both block conditions, stimulation consisted of 12 pure sinusoidal tones between 261.63 and 932.33Hz. They corresponded to musical tones C4, D4, E4, F#4, G#4, A#4, C5, D5, E5, F#5, G#5, and A#5 (261.63 Hz, 293.66 Hz, 329.63 Hz, 369.99 Hz, 415.30 Hz, 466.16Hz, 523.25 Hz, 587.33 Hz, 659.26 Hz, 739.99 Hz, 830.61 Hz, 932.33 Hz). From these 12 tones, four triplets, i.e., three tone sequences, were created. For better perceptibility, the tones within one tone triplet never spanned more than one octave.

In the structured condition, the same four triplets repeated over the course of a stimulation block, fitting with the experiment's original goal of study statistical learning (1). For each of the four triplet sequences, three different permutations were created, counterbalanced across participants, to control for the position of each individual tone within the triplet, i.e., each tone occurred in each triplet position —first, second or third — across the three permutations. In the random condition, the triplets were pseudo-randomly ordered. In both cases, each tone occurred equally often and there were neither consecutively repeating tones nor triplets. Each stimulation block consisted of 2400 tones, in which each tone was 300ms in duration with an interstimulus interval of 33ms, leading to a presentation period of 333 ms (i.e., repetition frequency = 3.0 Hz). This resulted in a total duration of 13.32 minutes per stimulation block. Each fetus was exposed to the random condition first and the structured condition second.

The auditory stimulation was produced by a loudspeaker outside of the shielded room and conducted through air-filled tubes into the shielded room. The tubes ended in an air-filled balloon (King Systems Corporation, Noblesville, USA) which was placed on the maternal abdomen. Stimuli were presented with an intensity of 95dB measured at the maternal abdomen. As the sound is attenuated through the maternal abdomen by about 30dB (2), the sound intensity which reaches the fetus is approximately 65dB.

Parallel experiments were also conducted using the same stimuli in a separate pool of infant and adult experiments. Infant data were not analyzed here due to the small sample size following data cleaning (n = 12);moreover, this infant sample was not studied before birth, unlike Experiment 2, which included longitudinal data before and after birth. Adult data were previously published (1, 3) in the context of the original study aims focused on statistical learning and are not analyzed here, as they are not relevant for our hypotheses.

**Artifact reduction and preprocessing .** Preprocessing took place in in MATLAB (The MathWorks Inc., Natick, MA, USA) version R2022b (artifact reduction) and R2024b. Magnetocardiography (MCG) data also recorded by the SARA device were preprocessed to extract heart rate variability (HRV) parameters. The fully automated R-peak detection algorithm (FLORA) was used to detect maternal and fetal R-peaks from the heart signal (4). Cardiac artifacts from the maternal heart were removed with a modified version of the fully automated subtraction of heart activity (FAUNA) algorithm (5). Briefly, the FAUNA algorithm is an in-house preprocessing procedure to remove cardiac activity from fetal biomagnetic data using principal component analysis (PCA) and ridge regression. FAUNA first constructs a template of the cardiac signal using the output of the FLORA algorithm. Next, spatial filtering is implemented with PCA and ridge regression to subtract cardiac activity. In a final step, independent component analysis (ICA) with the Fast ICA algorithm (6) is used to remove any residual cardiac activity remaining after the above steps. In the original published version of FAUNA (5), 40% of ICA components ranked by cross-correlation coefficients were removed, as well as ICA components that reached a threshold of 1 when averaged over all detected R-peaks. In the modified version used here, ICA components were removed more selectively using the statistical significance of their cross-correlation coefficients to avoid over-cleaning of the data.

Although the SARA device records 156 channels, the fetal head is small in relation to the sensor array. Thus, the next step in the preprocessing workflow is to identify fetal cortical channels. Following artifact removal, a cluster of ten cortical channels was identified that maximized variance in the delta (1 - 4 Hz) band. Only delta frequencies were considered for this purpose, so as to avoid biasing channel selection by muscle activity or noise. All channels in the identified cluster were broadband filtered (0.5 - 150 Hz) using a 2nd order Butterworth filter and isolated for MEG analysis.

In later processing steps, narrow band filtering was applied to MEG data (1 - 10 Hz, finite impulse response, filter order = 1221) and all signals were z-scored to normalize amplitudes across subjects. To avoid habituation effects (7), only data from the first 6 minutes of each stimulus condition were used, as prior frequency tagging studies conducted with infant EEG suggest that only several minutes of data are sufficient (8). Although stimuli were presented in continuous streams, we epoched data into 4 second trials, as this length yielded a large number of trials (n = 90), while also giving a roughly comparable data length as in Experiment 2 after trial-averaged stimulus conditions were horizontally concatenated (Experiment 1: 8 seconds, Experiment 2: 9.6 seconds; see Fig. 1, main manuscript). Finally, data were mean averaged across trials and RMS averaged across channels.

#### Experiment 2

**Study population.** Experiment 2 was conducted with the original aim of investigating hierarchical learning in fetuses and newborn infants using the global prediction error, which has been identified as a neural marker of consciousness in adults (9).

***Fetuses.*** Recruitment of pregnant volunteers followed informed consent and used the same inclusion and exclusion criteria as in Experiment 1 to yield an initial sample of 60 participants. Of these 60 participants, 56 pregnant women successfully completed at least one fetal MEG recording. Furthermore, 19 participants gave data at a first followup visit, 14 participants gave data at a second followup visit, and 5 participants gave data a third followup visit, resulting in a total data yield of 81 fetal MEG recordings.

***Newborns.*** After birth, many women returned to the laboratory with their newborn infant for further MEG data collection. In total, 33 newborns gave data in Experiment 2, of which 27 successfully completed the MEG recording; each newborn only gave data once. From the sample of 27 complete neonatal MEG recordings, 20 neonatal MEG recordings passed a quality control check. We used data from all 20 newborns in this sample, 16 of whom also had usable fetal data included in our analysis.

##### Experimental details.

***Fetuses.*** Following informed consent, the fetal head position was determined using ultrasound both before and after fetal MEG recordings. Fetal MEG data were recorded using the SARA device during two experimental phases: an exposure phase, during which fetuses passively learned an auditory sequence (30 trials), and a test phase (180 trials), during which fetuses were presented with both the familiar sequence (75% of trials) and an unfamiliar sequence (25% of trials). Crucially for the original aims of the experiment — though incidental to our aims in the current work — auditory oddballs could occur both within a sequence (local deviant) or between sequences (global deviants). Besides the two experimental phases, the experiment also used two blocks. In Block A, the subject was exposed to sequences of identical tones [ssss], i.e., the fetus was trained to anticipate no deviants within a sequence. In Block B, the subject was exposed to sequences ending in an oddball [sssd], i.e., the fetus was trained to anticipate a deviant within each sequence.

Each sequence consisted of four separate auditory tones at just two unique auditory frequencies (500 and 750 Hz) presented for 200 ms with a 400 ms interstimulus interval measured from one tone's offset to the next tone's onset. Auditory stimuli were presented using an MEG-compatible sound balloon at 90 dB sound intensity measured from the maternal abdominal surface, resulting in an estimated sound intensity of 60 dB for the fetus. For further details, see Moser et al. (10).

***Newborns.*** Stimuli for newborns were the same as described above for fetuses. Neonatal MEG data were recorded using the same SARA device as was used for fetal MEG. For neonatal recordings, each newborn was placed in a cradle and positioned head-first toward the MEG sensor array, such that the right temple of the newborn's head rested on the sensors. An MEG-compatible and infant-friendly headphone was then placed on the left ear for auditory stimulus delivery. Auditory tones were presented with the same experimental design as described above for fetuses using a sound intensity of 65 dB.

##### Preprocessing and data cleaning.

***Fetuses.*** Preprocessing took place in MATLAB R2016b. The FLORA algorithm (4) was used to identify maternal and fetal heartbeats. We then removed cardiac activity from fetal MEG using the FAUNA algorithm as described in Sippel et al. (5). Due to the relatively small extent of the fetal head in relation to the sensor array, a subset of 10 channels were chosen for further analysis by pinpointing a cluster of channels with the highest amplitudes after removing artifacts. Next, we bandpass filtered the selected channels 1 - 10 Hz using a fourth-order Butterworth filter, and the filtered signals were RMS averaged across channels. Then, trials were epoched from 200 ms before the onset of the first tone in a sequence to 1000 ms after the offset of the fourth tone in a sequence. Although global standard sequences were threefold more common than global deviant sequences, an equal number of trials (46 each) were averaged in both cases to balance the SNR across both conditions. Finally, to account for differences in fetal head position across recordings, each MEG recording was normalized as a percentage according to the evoked response in a 50 - 350 ms window after the first tone. Specifically, this reference value was computed by averaging signals across all trials pooled across both the exposure and the test phase and then finding the local maximum of the trial-averaged signal within the 50 - 350 ms window.

***Newborns.*** Preprocessing of neonatal MEG data in Experiment 2 followed different steps. For each neonatal recording, the magnetocardiogram (MCG) signal was detected through either template matching or the Hilbert transform. The MCG signal was then be subtracted from the data using a signal space projection, as outlined in previous work by others (11–13). We then bandpass filtered the data 1 to 15 Hz (Butterworth filter). High amplitude artifacts were attenuated using an artifact block algorithm, as detailed in Mourad et al.(14). The threshold for artifact detection was determined based on the median plus or minus 4 standard deviations of the data across all channels. If this range exceeds 2 pT, the MEG recording was excluded due to excessive noise. To best identify channels containing brain activity, an earlier approach from Moser et al.(15) was adapted. This approach uses principal component analysis (PCA). The first 3 principal components were considered, and their loadings were sorted to extract the 5 most influential channels. The locations of these 5 channels were determined and checked for plausibility based on two criteria: 1) the mean distances between channels should be $< 10$ cm, and 2) the channel locations should be within the area where the newborn's head was positioned. If multiple valid locations were identified, the location corresponding to the principal component with the highest explained variance was used. Next, selected channels were spatially RMS averaged, epoched into trials, and averaged across trials. Unlike fetal data from Experiment 2, neonatal data were *not* normalized according to a local maximum of the 50 - 350 ms neural response; rather, data were z-scored to normalize amplitudes prior to the frequency tagging analysis.

#### Perinatal variables

Several perinatal variables were collected and later correlated with frequency tagging strength. These were 1) the gestational age of the fetus or newborn, measured in weeks, 2) the birth weight, as later reported by parents for fetuses, 3) the maternal body mass index (BMI), as reported by the mother-to-be with respect to the very beginning of her pregnancy, 4) the maternal age, as measured in years, 5) the sex of the fetus or newborn, as later reported by the parents, and 6) the heart rate variability (HRV) of the fetus or newborn, taken as an indirect measure of arousal. Specifically, the HRV was measured from R peaks in the fetal or neonatal MCG signal extracted from the recorded data. Normal-to-normal R-R intervals were then calculated. The standard deviation of these intervals is the SDNN (16), which was taken in its original units (i.e., without log transformation) as the measured value of HRV.

#### Frequency tagging analysis

All usable data were entered into the frequency tagging analysis following channel and trial-averaging. The frequency tagging and all subsequent analyses were carried out in MATLAB R2022a and R2025a. Because each MEG recording encompassed multiple stimulus conditions which all shared a common stimulation frequency, conditions were concatenated across the dimension of time. Although horizontal concatenation of data introduces time domain discontinuities which may increase frequency domain noise, in this case, we reasoned that the slight increase in noise is more than compensated for by the benefit of adding more oscillatory cycles to the time domain signal, thus enhancing our ability to detect frequency tagging. For Experiment 1, this concatenation was done as [random, structured] after epoching data into 4-sec trials time locked to the onset of every $13^{th}$ tone (i.e., 12 tones per trial) and averaging across the first 90 epochs corresponding to 6 minutes of data, i.e., slightly less than half the total duration of each condition. Epochs from the remainder of each condition were discarded, as previous EEG frequency tagging work in infants has shown that $\leq 5$ minutes of data is sufficient (8) and, moreover, we reasoned that data in the second half of the experiment were more likely to induce habituation (7). For Experiment 2, the concatenation was performed as [ssss, sssD, sssd, sssS] on trial-averaged epochs, where capital letters denote global deviants. All usable trials from the test phase were averaged for each condition in Experiment 2, as stimuli were grouped into sequences rather than a constant stream, making habituation less likely.

In cases where data corresponding to one condition or training block were not usable due to noise, data from the remaining condition or training block were repeated to allow a common data length across all recordings, thereby avoiding data loss (as the alternative would be to discard recordings with missing data). Because all conditions contained stimulation with the same repetition frequency, we do not anticipate that this substitution was consequential to our hypothesis about frequency tagging besides benefiting the analysis through an increase in SNR. In Experiment 1, this substitution followed either [random, random] or [structured, structured] and increased the data yield from n = 20 to n = 31 fetal recordings. In Experiment 2, this substitution followed either [ssss, sssD, ssss, sssD] (repetition of data from block A) or [sssd, sssS, sssd, sssS] (repetition of data from Block B) and increased the data yield from n = 41 fetal recordings to n = 81 fetal recordings and from n = 15 to n = 20 neonatal recordings.

Our frequency tagging analysis closely follows prior analyses in the published literature, including EEG frequency tagging studies in infants. As described in the main manuscript, this analysis entails averaging all MEG signal recordings in the time domain to optimize SNR and then computing the raw fast Fourier transform (FFT, 0.05 Hz frequency resolution) without any frequency smoothing or windowing, as the effect of interest is highly frequency-specific. We then evaluated the frequency tagging effect on the FFT amplitude spectrum using separate parametric and nonparametric approaches, both described in the main manuscript. The parametric approach follows from previous EEG frequency tagging studies (8, 17–19) by z-scoring the FFT amplitude at the frequency of interest using the mean and standard deviation of the 20 neighboring amplitudes from 10 adjacent frequency bins on each side, excluding those that are immediate neighbors to the frequency bin of interest. The resulting z-score is then converted to a one-tailed P-value to test the frequency tagging hypothesis.

On the other hand, the nonparametric approach is a novel test we developed as an alternative to the parametric approach; it destroys the temporal alignment between MEG signals, prior to group averaging, via a surrogate transformation [iterative amplitude adjusted Fourier transform with exact distribution or 'IAAFT1' algorithm (20)] with code from Lancaster et al. (21) which preserves the power spectrum and data distribution of each signal while scrambling the Fourier phases. Surrogate signals were randomly generated and group-averaged 1000 times to build a null distribution for the parameter of interest, the z-scored SNR [Z(SNR)] at the stimulation frequency. The null distribution then allows an empirical P-value to be computed when the true value of Z(SNR) is compared to the null distribution. If Z(SNR) falls completely above the range of the null distribution (as was the case in Experiment 2 fetal data, see Table 1 main manuscript), then the P-value is estimated as 1/(N+1), where N = 1000 surrogates.

Because the nonparametric approach has not previously been applied to test for frequency tagging, we validated this novel test using publicly available EEG frequency tagging data from infants, previously evaluated using the parametric approach (8). In brief, these data were collected during a visual stimulation paradigm involving face images interleaved at 1.2 Hz with a periodic train of other, non-face images. 15 infants passively attended to the stimuli during EEG while seated on their mother's lap or in a car seat. We derived P-values for each of 32 EEG channels according to the nonparametric approach at the frequency of interest (1.2 Hz) and compared these values to the parametric P-values in the publication by de Heering and Rossion (8). All P-values were $-log_{10}$ transformed. We qualitatively compared the spatial topography of P-values derived from both approaches and we also computed the Pearson correlation between P-values from all channels.

For data visualization purposes, we also used 151 Morlet wavelets [code adapted from Hipp et al. (22, 23)] evenly spaced between 0.5 and 8.0 Hz to compute the time-averaged MEG power spectral density with spectral smoothing. In the case of fetal data from Experiment 2, which were normalized as a percentage, the group-averaged signal was z-scored to ensure zero mean for compatibility with the wavelet transform.

#### Subsample-based power analysis

Because we only measured frequency tagging at the group level, our analysis does not yield an effect size measure. Thus, to facilitate independent replication of our results and guide the design of future studies of fetal MEG frequency tagging, we estimated statistical power using a subsample-based approach. We computed Z(SNR) from many differently sized subsamples of data. Specifically, for all possible sample sizes in Experiment 1 (n = 1 to n = 31 in step size increments of 1), we drew subsamples of data without replacement 100 times, resulting in a total of 3100 subsamplings. Due to the need to repeat testing for each subsample, we focused only on the Z-test approach for testing frequency tagging, as the approach based on surrogate data would have been computationally expensive (3100 subsamples x 1000 surrogate data generations). We drew subsamples only from data in Experiment 1 because this experiment included only one MEG recording per fetus, ensuring statistically independent samples.

#### Bootstrapped resampled correlations

Having run parametric and nonparametric significance tests to evaluate our main hypothesis, we next computed correlations between frequency tagging strength and several perinatal variables listed earlier (see "Perinatal variables"). Because grand-averaging eliminates between-subjects variance, we utilized a bootstrapped resampling procedure, and sex was transformed to a continuous scale based on the average value of each bootstrapped resample (0 = male, 1 = female; e.g., 0.33 indicates a 2:1 male:female ratio within a resample). For each dataset of fetal or neonatal MEG recordings in each experiment, 500 bootstrapped resamples of size n were drawn with replacement. Both the MEG signal and perinatal variables were then averaged within each bootstrapped resample, and the z-scored SNR [Z(SNR)] was computed from the averaged signal. We computed the Pearson correlation coefficient between each perinatal variable and the Z(SNR) using all 500 resamples. This procedure was repeated for a total of 1000 iterations. The median Pearson coefficient across all iterations was then used to estimate the correlation strength, and a parametric P-value was computed with df = 498 (corresponding to a sample size of 500). We also computed an empirical 95% confidence interval using 1000 bootstrapped iterations.

As with the nonparametric significance test described earlier, we developed the bootstrapped resampling correlation test described above to the meet the needs of our study. Because this is a novel approach to estimating correlation strength, we validated the bootstrapped resampling procedure on synthetic data to confirm that the type I error rate does not increase with the number of resamples. To this end, we generated 50 pink noise signals, which follow a powerlaw distribution of spectral energy similar to MEG, and a 3 Hz sinusoid was added to each signal with a random weighting. The phase of the sinusoid was constant across signals to preserve temporal alignment. We then implemented the bootstrapped resampling procedure as described above, sampling with replacement from signals and the corresponding vector of weights. The resampled signals were then time-domain averaged and FFT transformed, and Z(SNR) was computed at 3 Hz. Similarly, the resampled weights were averaged using the mean. This was repeated a variable number of times to test different numbers of iterations ranging from n = 25 to 500 in steps of 25. In each case, n averaged resamples of Z(SNR) were correlated with n averaged resamples of weights. To infer the type I error rate, the n weights were randomly shuffled to destroy any relationship with Z(SNR). To infer the type II error rate, the ordering of the n weights was unchanged to preserve its relationship with Z(SNR).

### Supplemental Results

#### Fast Fourier transform applied to fetal and maternal cardiac data

To inform our understanding of peaks seen in the fast Fourier transform (FFT) of each group-averaged signal (Fig. 1, main manuscript), we applied the FFT to cardiac data to understand how cardiac signals appear in the FFT amplitude spectrum. We used a biomagnetic recording obtained in Experiment 1 at 34 weeks gestation. Data from 151 channels were taken at two different stages of preprocessing which isolated the maternal and fetal magnetocardiaogram activity separately. In each case, we applied principal component analysis (PCA) to the 151-channel data and extracted the first principal component which accounted for 91.8% and 99.2% of the variance in fetal and maternal cardiac data, respectively. Using the initial 4 seconds of data in each component, we computed the FFT and plotted the amplitude spectrum which, in each case, revealed a peak at the fetal or maternal heart rate but also at harmonic frequencies. These harmonics were extremely prominent and often of higher amplitude than the fundamental frequency. Because we did not observe harmonics in the group-averaged MEG amplitude spectra (Fig. 1, main manuscript), it is unlikely that any peaks therein were the result of cardiac artifacts.

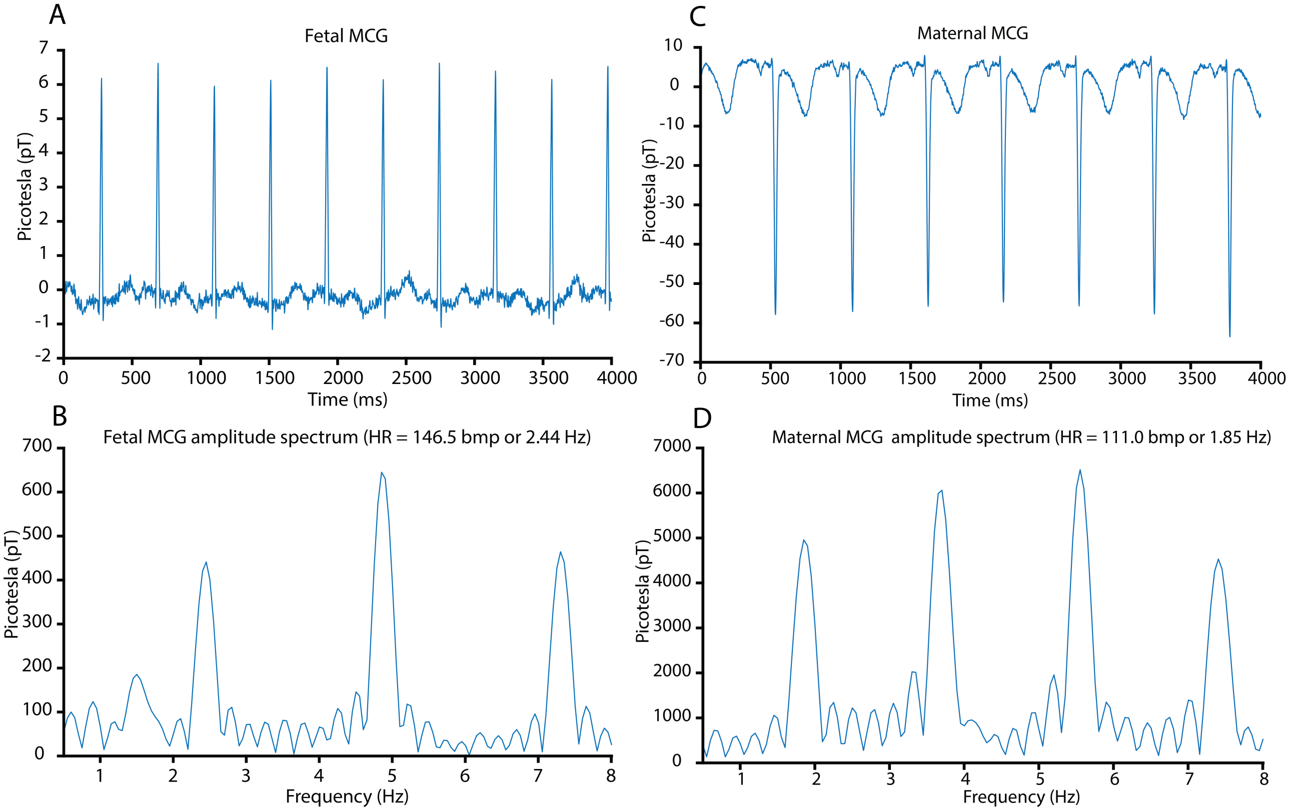


**Supplemental Figure 1** When the fetal magnetocardiogram (MCG) signal (A) is Fourier transformed using the FFT algorithm (fast Fourier transform), the resulting amplitude spectrum (B) shows multiple harmonics of the fundamental frequency corresponding to the fetal heart rate. The same results can be seen using the maternal MCG signal (C) and its FFT amplitude spectrum (D) with multiple harmonics.

#### Validation of the nonparametric frequency tagging approach

Using previously published infant EEG frequency tagging data (8), we showed that nonparametric and parametric approaches for testing a frequency tagging effect yield highly similar results (Fig. S1), with slight deviations that would be expected if each approach tested subtly different aspects of frequency tagging (see "Results and Discussion: Evidence of MEG frequency tagging in fetuses and newborns" in the main manuscript). To visualize the similarity of each approach, all P-values were $-log_{10}$ transformed and converted to topographic scalp maps (Fig. S1A,B). Both topographies show similar patterns of leftward dominance accompanied by a local maximum over the scalp vertex. Next, we computed the Pearson correlation coefficient of $-log_{10}(P)$ from both approaches (Fig. S1c) and found a very strong, positive correlation (r = 0.81, P = 2.5e-8). Several channels showed statistically significant ($\alpha = 0.05$, uncorrected) frequency tagging effects according to both approaches: F7, CP1, P7, O1, P8, and Cz (green area, Fig. S1C).

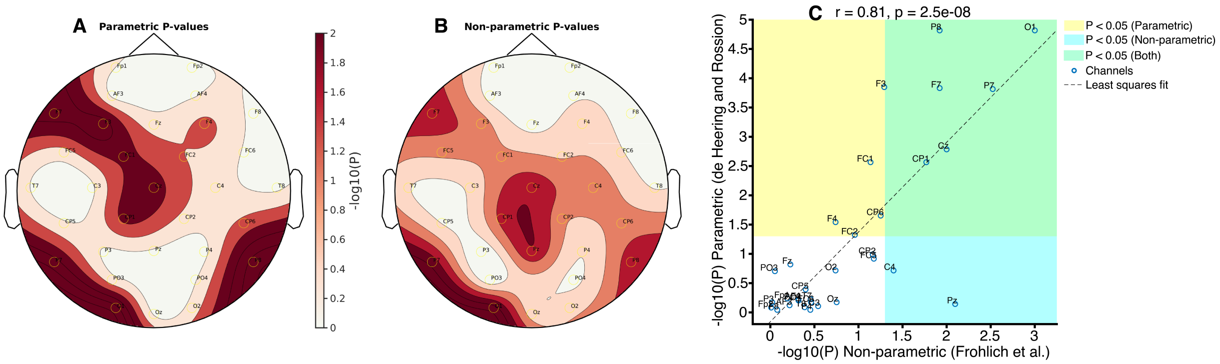


**Supplemental Figure 2** Panels (A) and (B) display P-values transformed as $-log_{10}(P)$, with similar scalp topographies in each case (e.g., local maxima over the scalp vertex, left parieto-occipital scalp, left frontal scalp, etc.). We then correlated the values transformed as $-log_{10}(P)$ from both the parametric and non-parametric approach and found a very strong linear relationship (C) as measured using the Pearson coefficient (r = 0.81)

#### Subsample-based statistical power analysis

Using a Monte Carlo subsampling approach, we determined that n = 13, 23, and 26 fetal MEG recordings are needed to detect a significant ($\alpha = 0.05$) frequency tagging effect (one-tailed Z-test) with 50%, 80%, and 90% statistical power, respectively, in Experiment 1 (see Fig. S3 below). For context, the sample size needed to design a fetal MEG frequency tagging study with 80% statistical power (n = 23) is nearly the median number of participants (n = 24) in general neuroimaging studies published by top journals in 2018 (24). Because fetal recordings were not independent samples in Experiment 2 (i.e., some pregnant women visited the laboratory multiple times to give longitudinal data), we did not attempt the same subsampling analysis using fetal data from Experiment 2.

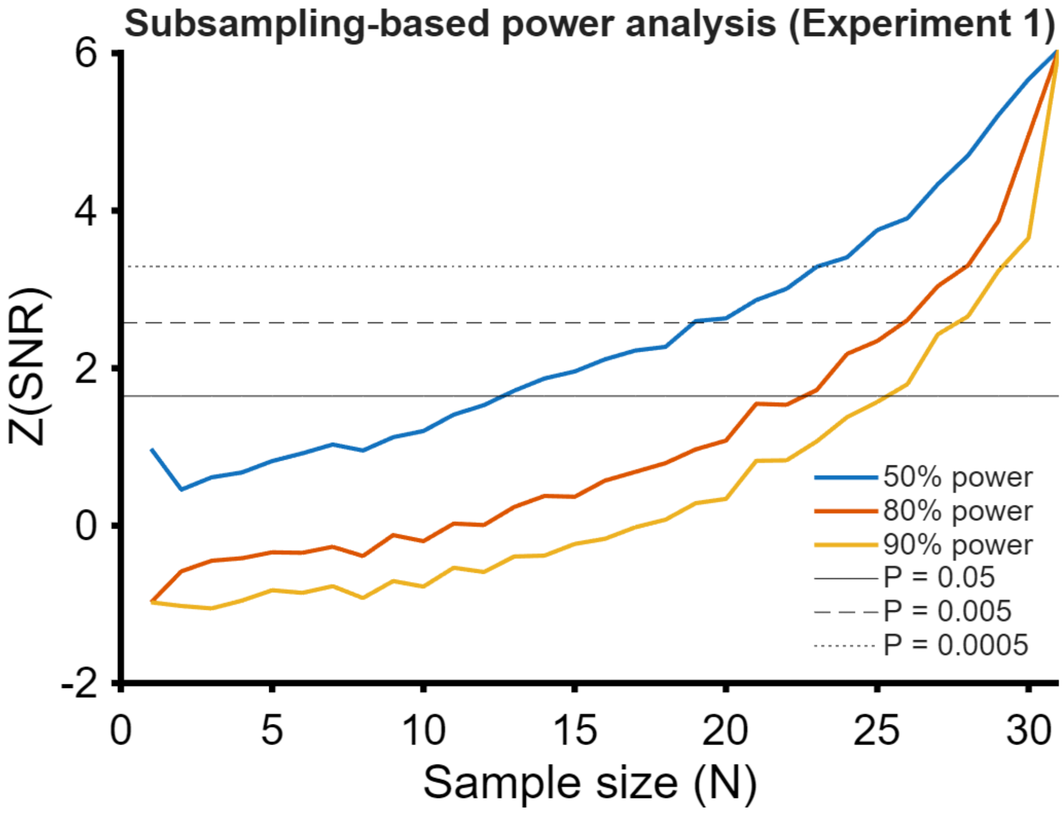


**Supplemental Figure 3** To estimate statical power, Monte Carlo subsampling was used with 100 subsampling for every possible sample size. We focused on Experiment 1, as this experiment only contained one fetal MEG recording per subject, i.e., each recording could be treated as an independent subject.

#### Spectrally smoothed MEG power spectra from Morlet wavelet transform

To infer frequency tagging, our analysis used the raw FFT amplitude spectra computed from group-averaged MEG signals (Fig. 1, main manuscript). While this approach is appropriate and necessary to detect frequency tagging (8, 17–19), we anticipate that many readers will be unfamiliar with seeing the raw FFT amplitude spectrum and more familiar with seeing the power spectrum (i.e., the amplitude spectrum squared) and, moreover, spectral smoothing (e.g., using a windowed or tapered approach). Thus, to contextualize the results in Fig. 1, we have provided the MEG power spectrum dervied from a Morlet wavelet transform with spectral smoothing below in Fig. S4 for each dataset. Note that while spectra displayed here are less noisy than the corresponding spectra in Fig. 1 of the main manuscript, frequency peaks are masked by smoothing across frequencies.

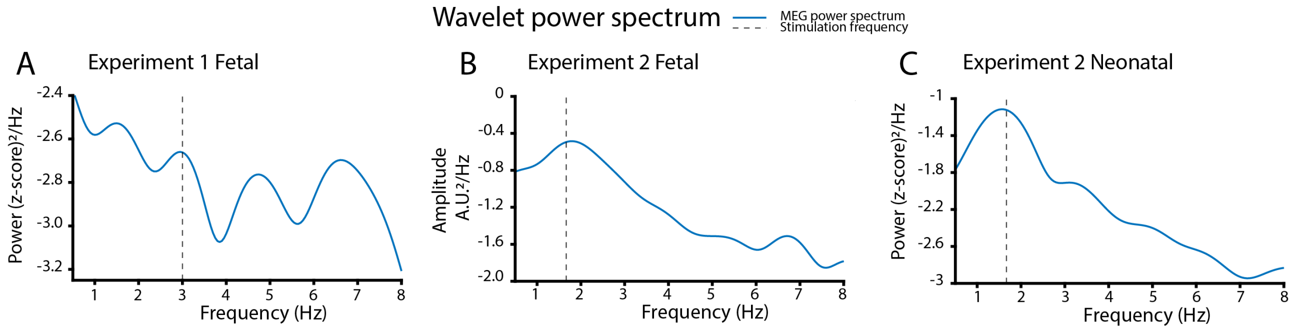


**Supplemental Figure 4** The stimulation frequency is marked with a vertical dashed line. A.U. = arbitrary units

#### Validation of the bootstrapped resampling correlation approach

Given the large number of resamples employed in our correlation analysis, we used simulated data to confirm that the number of resamples did not affect the type I error rate (i.e., false positive rate). To recap the approach described earlier (Supplemental Methods: Bootstrapped resampled correlations), we generated synthetic data for 50 simulated subjects using pink noise signals (i.e., power-law noise with $\alpha = 2$) with an added 3 Hz sinusoid. For each simulated subject, the sinusoid was weighted differently according to random coefficients with a uniform distribution, yielding a different SNR in all 50 cases. We then used our bootstrapping procedure described earlier to derive correlations between Z(SNR) at 3 Hz and the coefficients. 2000 bootstrapped resampling trials were used with a bootstrap size varying between n = 10 and n = 500 in incremental steps of 10. The type I error rate was estimated by correlating Z(SNR) with randomly shuffled coefficients and noting the number of false positives at $\alpha = 0.05$. Next, the type II error rate was estimated by correlating Z(SNR) with the unshuffled coefficients and noting the number of false negatives at $\alpha = 0.05$.

The Type I error rate remained nearly constant between 25 and 500 resamples (min = 0.038, max = 0.064, median = 0.050; see Fig. S2a). Moreover, the type II error rate fell monotonically following an exponential decay curve from 0.9 to 0 in the domain of 25 to 500 resamples (Fig. S2b). These results suggest that increasing the number of resamples in our bootstrapped correlation analysis only decreases the type II error rate (proportion of false negatives) without increasing the type I error rate (proportion of false positives). We concluded that a larger bootstrap size improves the type II error rate at no expense to the type I error rate.

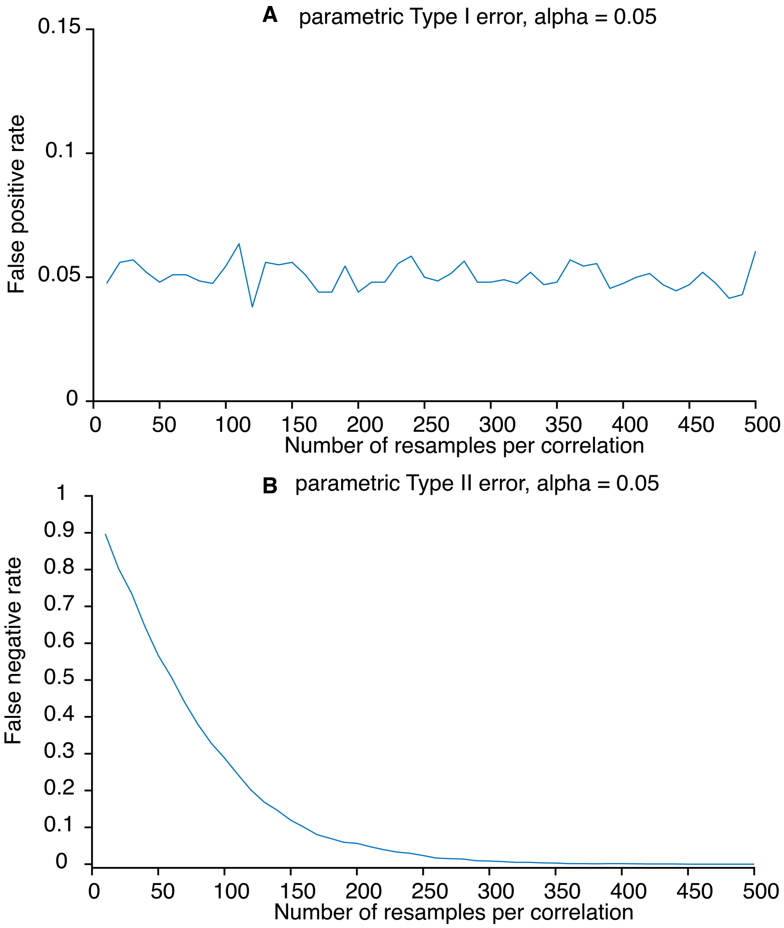


**Supplemental Figure 5** In simulated data, the type I error rate remains stable regardless of the bootstrap size (A). Conversely, the type II error rate decreases monotonically as the bootstrap size decreases.

#### Correlational analysis with bootstrapped resampling

We next explored whether frequency tagging strength varies with perinatal variables. Given the large number of resamples employed in our correlation analysis, we first used simulated data to confirm that the number of resamples did not affect the type I error rate, which remained nearly constant between 25 and 500 resamples (Fig. S3). In fetal and newborn data, bootstrapped resampling revealed a number of significant correlations between Z(SNR) and perinatal variables. None of the correlations tested were significant in both experiments, except for maternal age, where the sign of the Pearson coefficient disagreed between fetal experiments (Table 1). However, within Experiment 2, a strongly significant ($P_{FDR} < 0.005$) positive correlation between maternal age and Z(SNR) replicated between fetal (r = 0.14) and newborn (r = 0.18) data. This relationship is unlikely to be mediated by maturational changes in body weight distribution, as no significant correlation was detected between maternal body mass index (BMI) and Z(SNR) in fetal data from Experiment 2; however, note a significant relationship in Experiment 1 (Table 1).

Next, we examined correlations at the control frequency to infer whether the relationship with maternal age was specific to the experimental frequency. While the control frequency also yielded a significant correlation between Z(SNR) and maternal age in all three datasets (Experiment 1 fetal, Experiment 2 fetal, and Experiment 2 neonatal). However, in each case, the Pearson coefficients were significantly different between the experimental and control frequencies according to the Fischer R to Z transformation ($P < 0.05$, uncorrected). The foregoing evidence suggests unique relationship between maternal age and Z(SNR) at the experimental frequency, though this relationship differed between experiments.

Even within the common stimulation paradigm of Experiment 2, some correlations observed in neonatal data were not observed in fetal data, including, most notably, a strongly significant positive correlation with gestational age in newborns (Table S1). Given, however, the many differences between newborns and fetuses (9, 25, 26), it is plausible that birth changes relationships between Z(SNR) and perinatal variables such that they are not reproducible from fetus to infant, even longitudinally.

### Supplemental Tables

| Test | $R_{median}$ | CI lower bound | CI upper bound | $P_{raw}$ | $P_{FDR}$ | Data | Frequency | Significance |
|---|---|---|---|---|---|---|---|---|
| age | -0.203 | -0.274 | -0.122 | 4.79e-06 | 2.15e-05 | Exp. 1 fetal | Experimental | *** |
| birth weight | 0.00685 | -0.0847 | 0.093 | 0.879 | 0.879 | Exp. 1 fetal | Experimental | |
| BMI | 0.134 | 0.0515 | 0.223 | 0.00278 | 0.00483 | Exp. 1 fetal | Experimental | ** |
| maternal age | -0.191 | -0.277 | -0.11 | 1.69e-05 | 6.07e-05 | Exp. 1 fetal | Experimental | *** |
| Sex | -0.14 | -0.226 | -0.0582 | 0.00171 | 0.00342 | Exp. 1 fetal | Experimental | ** |
| HRV | 0.0135 | -0.0713 | 0.0939 | 0.763 | 0.808 | Exp. 1 fetal | Experimental | |
| age | -0.0647 | -0.151 | 0.0224 | 0.149 | 0.206 | Exp. 2 fetal | Experimental | |
| birth weight | 0.225 | 0.141 | 0.308 | 3.62e-07 | 2.17e-06 | Exp. 2 fetal | Experimental | *** |
| BMI | -0.0178 | -0.109 | 0.0766 | 0.691 | 0.808 | Exp. 2 fetal | Experimental | |
| maternal age | 0.142 | 0.0536 | 0.23 | 0.00148 | 0.00332 | Exp. 2 fetal | Experimental | ** |
| Sex | -0.0491 | -0.139 | 0.0431 | 0.273 | 0.351 | Exp. 2 fetal | Experimental | |
| HRV | -0.088 | -0.172 | 0.00271 | 0.0493 | 0.0739 | Exp. 2 fetal | Experimental | |
| age | 0.275 | 0.19 | 0.355 | 3.92e-10 | 3.53e-09 | Exp. 2 newborn | Experimental | *** |
| birth weight | -0.0142 | -0.101 | 0.067 | 0.751 | 0.808 | Exp. 2 newborn | Experimental | |
| BMI | 0.308 | 0.23 | 0.386 | 1.78e-12 | 3.21e-11 | Exp. 2 newborn | Experimental | *** |
| maternal age | 0.178 | 0.0926 | 0.264 | 6.04e-05 | 0.000181 | Exp. 2 newborn | Experimental | *** |
| Sex | 0.133 | 0.0524 | 0.219 | 0.00295 | 0.00483 | Exp. 2 newborn | Experimental | ** |
| HRV | 0.145 | 0.0541 | 0.233 | 0.00117 | 0.003 | Exp. 2 newborn | Experimental | ** |
| age | -0.000504 | -0.0879 | 0.0785 | 0.991 | 0.997 | Exp. 1 fetal | Control | |
| birth weight | -0.156 | -0.246 | -0.0697 | 0.000448 | 0.00134 | Exp. 1 fetal | Control | ** |
| BMI | 0.0987 | 0.0112 | 0.184 | 0.0273 | 0.0446 | Exp. 1 fetal | Control | * |
| maternal age | -0.0652 | -0.149 | 0.0232 | 0.146 | 0.187 | Exp. 1 fetal | Control | |
| Sex | 0.189 | 0.108 | 0.268 | 2.11e-05 | 9.51e-05 | Exp. 1 fetal | Control | *** |
| HRV | -0.216 | -0.297 | -0.132 | 1.04e-06 | 9.38e-06 | Exp. 1 fetal | Control | *** |
| age | 0.133 | 0.0546 | 0.218 | 0.0028 | 0.00629 | Exp. 2 fetal | Control | * |
| birth weight | -0.0669 | -0.149 | 0.0199 | 0.135 | 0.187 | Exp. 2 fetal | Control | |
| BMI | -0.0933 | -0.182 | -0.00444 | 0.037 | 0.0555 | Exp. 2 fetal | Control | |
| maternal age | -0.103 | -0.187 | -0.0121 | 0.0215 | 0.0387 | Exp. 2 fetal | Control | * |
| Sex | 0.000179 | -0.0869 | 0.0906 | 0.997 | 0.997 | Exp. 2 fetal | Control | |
| HRV | 0.241 | 0.155 | 0.322 | 4.74e-08 | 8.54e-07 | Exp. 2 fetal | Control | *** |
| age | -0.0182 | -0.102 | 0.07 | 0.685 | 0.823 | Exp. 2 newborn | Control | |
| birth weight | 0.206 | 0.119 | 0.299 | 3.48e-06 | 2.09e-05 | Exp. 2 newborn | Control | *** |
| BMI | 0.136 | 0.0459 | 0.225 | 0.00235 | 0.00604 | Exp. 2 newborn | Control | * |
| maternal age | -0.175 | -0.261 | -0.0838 | 8.23e-05 | 0.000296 | Exp. 2 newborn | Control | *** |
| Sex | -0.13 | -0.219 | -0.0379 | 0.00351 | 0.00702 | Exp. 2 newborn | Control | * |
| HRV | 0.00676 | -0.0766 | 0.0979 | 0.88 | 0.99 | Exp. 2 newborn | Control | |

**Table 1. Statistical results of correlational analysis with bootstrapped resampling. CI = 95% confidence interval, Exp. = experiment**

$P_{FDR}$ **< 0.05 ***
$P_{FDR}$ **< 0.005 ****
$P_{FDR}$ **< 0.0005 *****

### Data availability

Data used in Experiment 1 of this study will be uploaded to a public repository at the time of publication. Data used in Experiment 2 of this study, originally published in Moser et al. 2021 (10), are already publicly archived and available through Zenodo: https://zenodo.org/record/4541463#.Y0a-iExByHt.
